# GeTFEP: A general transfer free energy profile of transmembrane proteins

**DOI:** 10.1101/191650

**Authors:** Wei Tian, Hammad Naveed, Meishan Lin, Jie Liang

**Affiliations:** Department of Bioengineering, University of Illinois at Chicago, Chicago, IL 60607; Department of Computer Science, National University of Computer & Emerging Sciences (NUCES-FAST), Islamabad, ICT 44000

## Abstract

Free energy of transferring amino acid side–chains from aqueous environment into lipid bilayers, known as transfer free energy (TFE), provides important information on the thermodynamic stability of membrane proteins. In this study, we derived a TFE profile named General Transfer Free Energy Profile (GeTFEP) based on computation of the TFEs of 58 *β*–barrel membrane proteins (*β*MPs). The GeTFEP agrees well with experimentally measured and computationally derived TFEs. Analysis based on the GeTFEP shows that residues in different regions of the TM segments of *β*MPs have different roles during the membrane insertion process. Results further reveal the importance of the sequence pattern of transmembrane strands in stabilizing *β*MPs in the membrane environment. In addition, we show that GeTFEP can be used to predict the positioning and the orientation of *β*MPs in the membrane. We also show that GeTFEP can be used to identify structurally or functionally important amino acid residue sites of *β*MPs. Furthermore, the TM segments of *α*–helical membrane proteins can be accurately predicted with GeTFEP, suggesting that the GeTFEP captures fundamental thermodynamic properties of amino acid residues inside membrane, and is of general applicability in studying membrane protein.

## 1 Introduction

Membrane proteins play important roles in cellular metabolism, signaling regulation, and intercellular interactions.^1^ Knowledge of the thermodynamic stability of membrane proteins is essential for understanding their folding behavior and their structure–function relationship.^2–5^ A widely used measure to estimate the stabilities of membrane proteins is the transfer free energies (TFEs), which quantify the free energies of transferring amino acid residues from aqueous environment into lipid bilayers.^6–11^

Often called hydrophobicity scales, transfer free energies have been measured experimentally based on several model systems. The Wimley–White whole residue scale (WW–scale) measures TFEs of residue partitioning between water and octanol using a set of peptides as the host of amino acids.^8^ The biological scale (Bio–scale) of Hessa *et al*. measures the free energies required to transfer residues in polypeptides into the ER membrane through the translocon machinery.^9^ The Moon–Fleming whole protein scale (MF–scale) measures TFEs of residues from water to the membrane core in the context of a whole *β*–barrel membrane protein (*β*MP).^10^ These experimentally obtained hydrophobicity scales have leaded to improved understanding of the structures and functions of membrane proteins^12^ and have been used in prediction of transmembrane (TM) segments of membrane proteins.^13^

However, experimental measurement of TFEs is technically challenging, cumbersome, and costly.^14,15^ Complementing experimentally measured transfer free energies, several hydrophobicity scales have been derived computationally, which can aid in our understanding of the governing principles of membrane protein folding.^2,4,16^ The *E*_*Z*_*α* and *E*_*Z*_*β* empirical potentials are knowledge–based hydrophobicity scales. They have been successfully applied in predicting the positioning of membrane proteins in the lipid bilayer, in discriminating side– chain decoys, and in identifying protein–lipid interfaces.^17,18^ However, these scales obtained from statistical analysis do not consider the physical interactions either between residues from neighboring helices/strands or within the same helix/strand, which are known to be important for membrane protein folding.^19,20^ There have also been studies based on molecular dynamics (MD) simulations to calculate TFEs,^21–23^ although the choice of the reference state before membrane insertion remains a challenging task.^22^

Another method of deriving TFEs computationally was developed for *β*MPs recently.^11^ This method is based on a statical mechanical model in a discretized conformational space. It incorporates both intra– and inter–strand interactions in the TM segments of the proteins. It can be used to calculate TFEs of any lipid–facing residue in the TM segment of a *β*MP, as long as the number of TM strands of the protein is no more than 12. The computed TFE scale (OmpLA scale) is in excellent agreement with the MF–scale with a Pearson correlation coefficient *r* = 0.90. This scale has been applied successfully to explain how the functional fold and topology of the *β*MP are determined by the asymmetry of both the Gram–negative bacterial outer membrane and the TM residues.^11^ A further algorithmic extension of this method has greatly reduced the computational cost, enabling the calculation of TFEs on all *β*MP known so far, regardless their sizes, with little loss of the accuracy.^24^

In this study, we use the new algorithm^24^ to compute the depth–dependent TFE profile of each *β*MP in a non–redundant set of 58 *β*MPs. After examining their overall patterns, we found that there exists a general TFE profile applicable to all *β*MPs, which we call the <u>Ge</u>neral <u>T</u>ransfer <u>F</u>ree <u>E</u>nergy <u>P</u>rofile (GeTFEP). The GeTFEP agrees well with previously measured and computed TFEs. Analysis based on GeTFEP shows that residues in different regions of the TM segment have different roles during the membrane insertion process. Our results further reveal the importance of the sequence pattern of TM segments in stabilizing *β*MPs in the membrane environment. In addition, we also show that GeTFEP can be used to predict positioning and orientation of *β*MPs when embedded in the membrane, with overall results in good agreement with experimental data. Furthermore, we show that the GeTFEP can be used to locate structurally or functionally important sites of *β*MPs. In addition, TM segments of *α*–helical membrane proteins can also be accurately predicted using the GeTFEP, suggesting that the GeTFEP captures fundamental thermodynamic properties of amino acid residues inside membrane, and has general applicability in studying membrane protein.

## 2 Results

### GeTFEP: <u>Ge</u>neral <u>T</u>ransfer <u>F</u>ree <u>E</u>nergy <u>P</u>rofile

#### Computation of TFE profiles of *β*MPs

Using the methods described in Ref [^24^], we calculate the depth–dependent TFE profiles for each *β*MP in a non–redundant set of 58 *β*MPs. The proteins in this set have *≤* 30% pairwise sequence similarity. Briefly, for each *β*MP, we substituted each lipid–facing residue in the TM region to the other 19 amino acids. We calculated the TFEs of each amino acid substitution using Ala as the reference. The TFE profile of the protein was then obtained by taking average of the TFE values of the same amino acid type at the same depth position in the membrane. As an example, Fig. 1 shows the computed TFE profile of the protein LptD, the largest *β*MP with known structure (PDB ID: 4q35).

**Figure 1.**
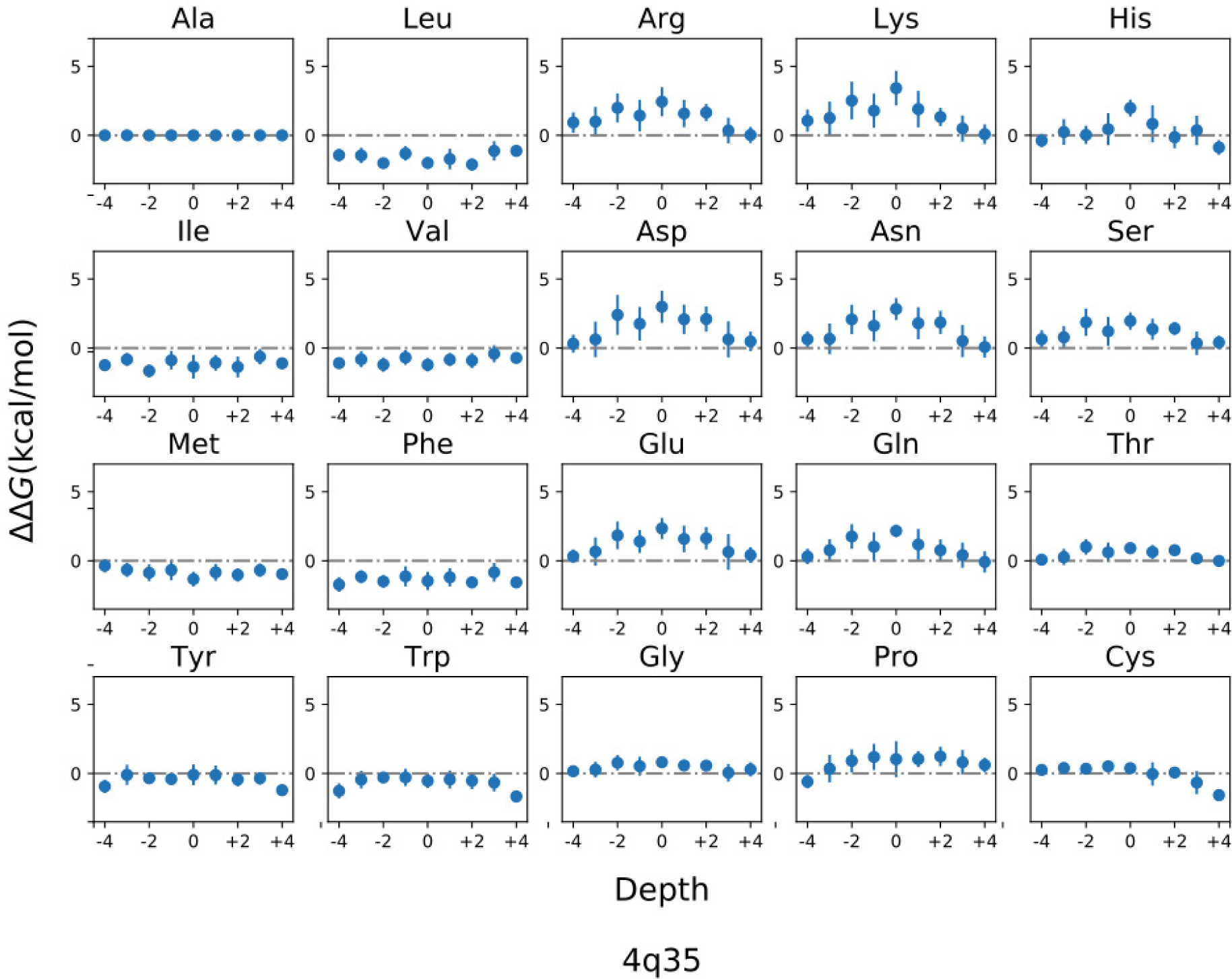
The TFE profile of the LptD protein. Each subfigure shows the calculated free energies (*y*–axis) of a specific amino acid type when transferred to certain depth position of the membrane (*x*–axis). The depth position of the membrane is discretized and indexed from −4 to +4, starting from the periplasmic side to the extracellular side of the outer membrane. Depth position 0 is at the mid–plane of the membrane.

#### Derivation of GeTFEP

Although the 58 *β*MPs are in different oligomerization states, have different sizes (strand numbers) of TM segments, and come from different organisms, their TFE profiles are remarkably similar. Results of clustering their profiles show that the 58 *β*MPs can be grouped into only one group (with 56 *β*MPs) and two outliers: *α*– and *γ*–hemolysins (PDB ID: 7ahl and 3b07). Details of the clustering method can be found in the supporting information.

Unlike the other *β*MPs, the TM regions of both *α*– and *γ*–hemolysins are formed by repeated *β*–hairpin (Fig. S2B), which make their TFE profiles highly sensitive to the composition of the *β*–hairpin and the local interactions of residues within the hairpin (Fig. S2 C and D). Accordingly, we further investigate whether *α*– and *γ*–hemolysins have truly different thermodynamic properties than *β*MPs, or their outlier status is due to the special architecture of repeated *β*–hairpins.

We first computed the TFE profiles of artificially generated hemolysin–like *β*MPs constructed by repeating each *β*–hairpin in our *β*MP set. Altogether, we computed TFE profiles for 778 artificial hemolysin–like *β*MPs. We then sampled from these profiles with replacement, and computed the distribution of the distance from each sampled profile to the average profile of all sampled artificial *β*MPs. The distances from the TFE profiles of both *α*– and *γ*– hemolysins to the average profile are at the 80th percentile in the distance distribution (Fig. 2B), indicating that *α*– and *γ*–hemolysins are not fundamentally different in their thermodynamic properties from other *β*MPs. Therefore, we conclude that a general transfer free energy profile exists and is applicable to all *β*MPs, including *α*– and *γ*–hemolysins. We derive the <u>Ge</u>neral <u>T</u>ransfer <u>F</u>ree <u>E</u>nergy <u>P</u>rofile (GeTFEP) by averaging the TFEs of a specific amino acid at the same lipid bilayer depth position for all 58 *β*MPs (Fig. 2C).

**Figure 2.**
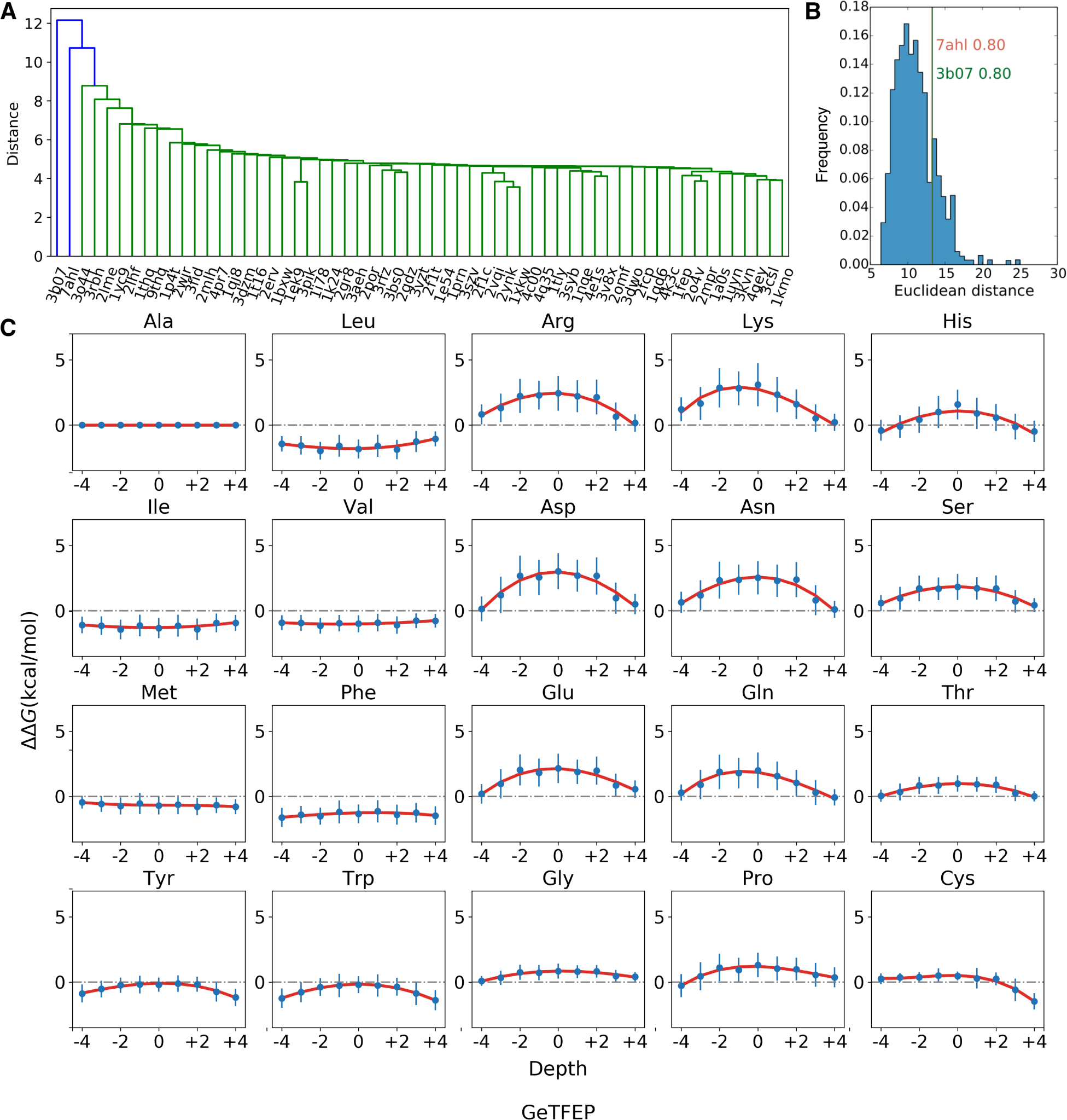
Derivation of the GeTFEP. **A.** Results of hierarchical clustering shows that all *β*MPs in the dataset can be group into one cluster, except *α*– and *γ*–hemolysins (7ahl and 3b07). **B.** The distribution of distance between the sampled TFE profiles of the artificially constructed hemolysin–like *β*MPs and their average TFE profile. The distances of both *α*– and *γ*–hemolysins are at the 80th percentile of the distribution. **C.** The General Transfer Free Energy Profile (GeTFEP) of each residues (blue), and the corresponding curves fitted by 3rd degree polynomials (red).

#### Comparison with other hydrophobicity scales

We then examine how GeTFEP compares with other hydrophobicity scales. Since most experimentally measured scales are not depth–dependent, we first compare the scale of the TFEs at the hydrocarbon core position of depth 0 in the GeTFEP with other hydrophobicity scales. We refer this hydrophobicity scale as the mid–GeTFEP scale. The mid–GeTFEP scale correlates well with the experimentally measured hydrophobicity scales, having Pearson correlation coefficients *r* = 0.83 with the WW–scale, and *r* = 0.92 with the Bio–scale. It also correlates well with the computational *β*MP OmpLA scale,^11,24^ with *r* = 0.90 (Fig. S3). When compared with the experimentally measured MF–scale of the *β*MP OmpLA mid– GeTFEP has a correlation of *r* = 0.87. One noticeable difference between mid–GeTFEP and the MF–scale is that the TFE value of His is less unfavorable in mid–GeTFEP (Fig. 3A).

**Figure 3.**
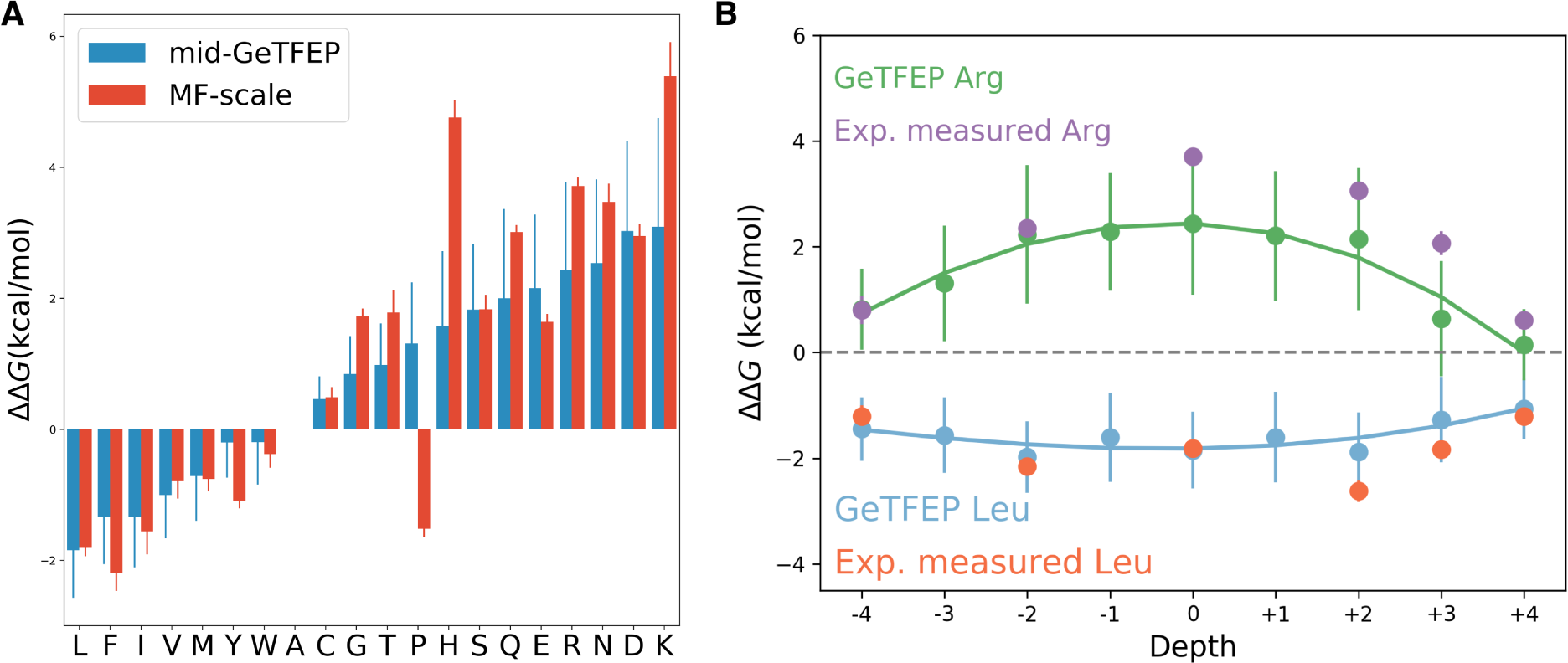
Comparison between the GeTFEP and the experimentally measured MF–scale. **A.** The mid–GeTFEP scale agrees well with the MF–scale, except Pro and His. **B.** The depth–dependent TFEs of Arg and Leu of GeTFEP also agree well with the experimental measurements.^10^

This is expected since the MF–scale was measured in acidic condition at pH=3.8, where His was fully protonated.^10^ The different value in mid–GeTFEP likely reflects the property of His in physiological conditions of the outer membrane.

Another notable difference is Pro. It is found that Pro is unfavorable in the membrane environment according to the mid–GeTFEP scale, while it is found to be favorable according to the MF–scale (Fig. 3A). Pro tends to disrupt the structures of both *α*–helix and *β*–sheet, and is thermodynamically unfavorable in the non–polar core of the membrane.^25^ The value of Pro in the GetFEP–mid scale reflects the general situation.

We then examined the depth–dependency of the GeTFEP of Arg and Leu, whose experimental results are available.^10^ Their TFEs at different depth positions of the membrane are in good agreement with the experimentally measured values, with *r* = 0.87 for Arg and *r* = 0.75 for Leu (Fig. 3B), suggesting the GeTFEP captures the depth–dependency of TFEs of amino acids.

### Insertion of *β***MP into membrane**

#### *β*MP insertion as a thermodynamically driven spontaneous process

Upon synthesis in the cytoplasm, *β*MPs need to be transported across the periplasm and then folded into the outer membrane. As there is no energy source such as ATP in the periplasm, it was suggested that the free energies of *β*MP folding provide an adequate source to ensure successful periplasm translocation.^26^ A computational study showed that the TFE of lipid–facing residues of the hydrophobic core regions are indeed the main driving force for membrane insertion.^11^ Analysis also showed that lipid–facing residues in the TM regions of of *β*MPs have clear patterns of amino acid composition.^27^ However, it is still unclear whether the insertion of *β*MPs into the membrane is primarily due to the extensive property of the hydrophobicity of lipid–facing residues, or the specific pattern of amino acid composition also plays important roles.

To investigate this question, we employed a simplified *β*MP insertion model based on the concerted folding mechanism proposed in Ref [^28^]. We ignore the effects of non–TM loops and discretizes the insertion process into 17 steps (Fig. 4A). We take the position recorded in the widely–used Orientations of Proteins in Membranes (OPM) database^29^ as the fully inserted position of each *β*MP. This position is denoted as the reference positioon 0, and the other positions are indexed accordingly from *-*8 to +8. *β*MPs start the insertion process at position *-*8 from periplasmic side and become fully inserted into the membrane at position 0. From position 0 to +8, *β*MPs would translocate across the membrane. We assume that the stability of the TM region of a *β*MP can be approximated by summarizing TFEs of all lipid– facing residues in the membrane region. The stability of the *β*MP at each position was then calculated using the GeTFEP following this additive model. As an example, Fig. 4B shows stability of the protein OmpA (PDB ID: 1bxw) at different insertion positions. Overall, results of all *β*MPs show a funnel–like pattern of insertion energy (Fig. 4C). Most *β*MPs (52 of 58) have minimum free energy when they are fully inserted into membranes (position 0, Tables S1 and S2). The funnel–like pattern indicates that the insertion of *β*MPs into outer membranes is indeed a spontaneous process. *β*MPs become energetically trapped after being fully inserted.

**Figure 4.**
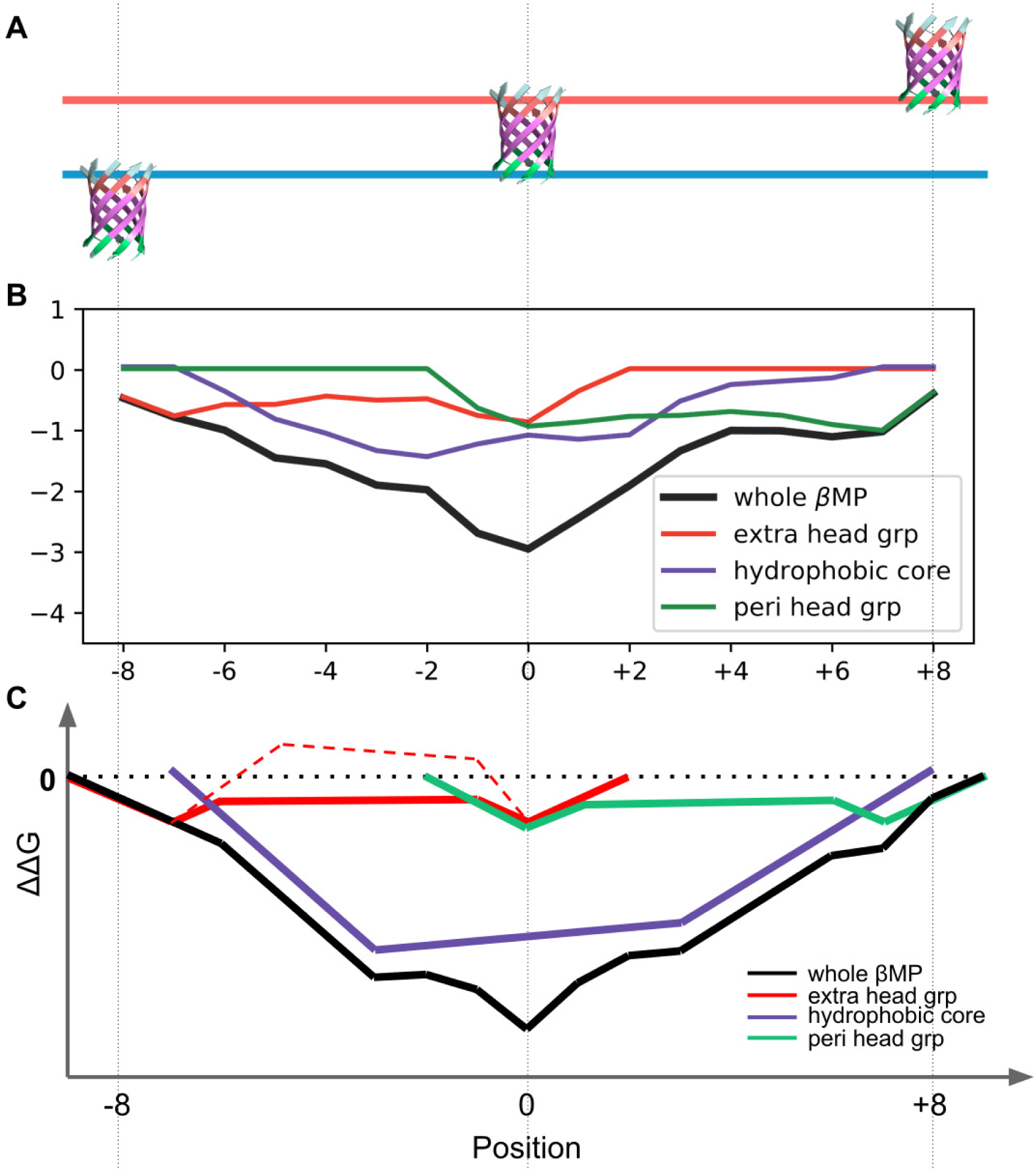
Illustration of the membrane insertion process of *β*MPs. **A.** The simplified *β*MP insertion model following Ref [^28^]. **B.** The computed insertion energies of the OmpA protein at different depth positions. Contribution of different regions are color coded. **C.** Illustration on how free energies change with the position of *β*MP in the membrane. The dashed red segments show that lipid–facing residues in extracellular head group region sometimes become energetically unfavorable.

#### Importance of patterns of TM lipid–facing residues in membrane insertion

We then examine if the funnel–like insertion energy pattern arises from the extensive property of the TFEs of the hydrophobic residues alone. We considered only the 52 *β*MPs whose minimum free energies are at the fully inserted position. We first shuffled the sequences of the *β*–strands within the TM segment of each *β*MP. While the side–chain direction as well as the interstrand hydrogen bond pairing at each residue position in *β*–strands are maintained, all TM residues are permuted. Each *β*MP is shuffled 2,000 times. We found that it is highly unfavorable to insert the shuffled *β*MPs into the membrane. This is expected, since the shuffling changes hydrophobicity of TM segments *β*MPs. Before the shuffling, the ionizable/polar residues were enriched among lumen–facing residues of *β*MPs, while lipid– facing residues were mostly apolar. After the shuffling, they were much evenly distributed.

We then investigate how insertion energy is affected if only the lipid–facing residues are shuffled. While the insertion of the shuffled *β*MPs remains energetically favorable (see Fig. S4 for an example), shuffled *β*MPs are less stable compared to the original *β*MPs at the fully inserted position for 50 out of 52 *β*MPs: The insertion energy for the shuffled *β*MPs is on average 6.36 kcal/mol higher (Table S1). In addition, the fully inserted position (position 0) is no longer the most stable position for 17.4% of the shuffled *β*MPs (Table S1). These results indicate that the locational patterns of lipid–facing residues^30^ in the TM region are optimized for *β*MPs to gain stability in the membrane environment.

#### Roles of residues in different TM regions during membrane insertion

The TM segment of a *β*MP can be divided into three regions, namely, the periplasmic headgroup region, the hydrophobic core region, and the extracellular headgroup region.^27^ We investigate how these regions contribute to the insertion energy of the *β*MP. We found that residues in the same regions across all 52 *β*MPs shared similar patterns in their insertion free energy profile (Fig. 4C), indicating that they play similar roles in the insertion process. Among these, lipid–facing residues of the extracellular headgroup region facilitate the initialization of the insertion process, as they are energetically favorable in the interfacial region on the periplasmic side (position −8 and −7). As insertion proceeds, these residues become less favorable and occasionally unfavorable when they become more embedded in the membrane. At this time, lipid–facing residues of the hydrophobic core region start to be inserted in the membrane, and strongly drive the insertion process (position −6 to −2). When lipid–facing residues of the extracellular headgroup region approach the interfacial region of the extracellular side, they become energetically favorable again. At the same time, lipid– facing residues of the periplasmic headgroup region become inserted (position −1 and 0), and the TFE of the whole *β*MP reaches its minimum at position 0.

Although lipid–facing residues of the hydrophobic core region are known to provide the main driving force for membrane insertion of *β*MPs,^11^ we found that the TFEs of hydrophobic core region do not reach their minimum when *β*MPs are fully inserted at position 0 for all 52 *β*MPs. Upon incorporation of contributions from other regions, the overall TFEs of the whole *β*MPs indeed reach the minimum at the fully inserted position. The “W” shape of the free energy curves of the two head group regions (the red and green curve in Fig. 4C) suggests that lipid–facing residues in these regions act like “energetic latches” to lock *β*MPs into their fully inserted position.

#### Prediction of *β*MP positioning and orientation in the membrane

GeTFEP can be used to predict positioning and orientation of *β*MPs in the membrane, similarly to previous studies.^17,18^ Here, the membrane is idealized as an infinite slab with a thickness of *h*. Each *β*MP is initially positioned in the membrane with its center of mass of the barrel domain at the midplane of the membrane and its barrel axis aligned with the normal direction (*z*–axis) of the membrane (Fig. 5A). The protein can be rotated around the *x*– and *y*–axes with angles *θ*_*x*_ and *θ*_*y*_, respectively. The two rotation angles together determine the tilt angle of the protein. The protein can also be translated with a displacement *d*_*z*_. This displacement and the membrane thickness determine the TM segment of the protein. When embedded in the membrane, the lipid–facing residues of the TM region and the loop residues are used to calculate the total energy of the *β*MP using the GeTFEP. As an example, Fig. 5B shows how rotation angles *θ*_*x*_ and *θ*_*y*_ affect the stability of the protein BtuB (PDB ID: 1nqe) when the displacement *d*_*z*_ and the membrane thickness *h* are fixed.

**Figure 5.**
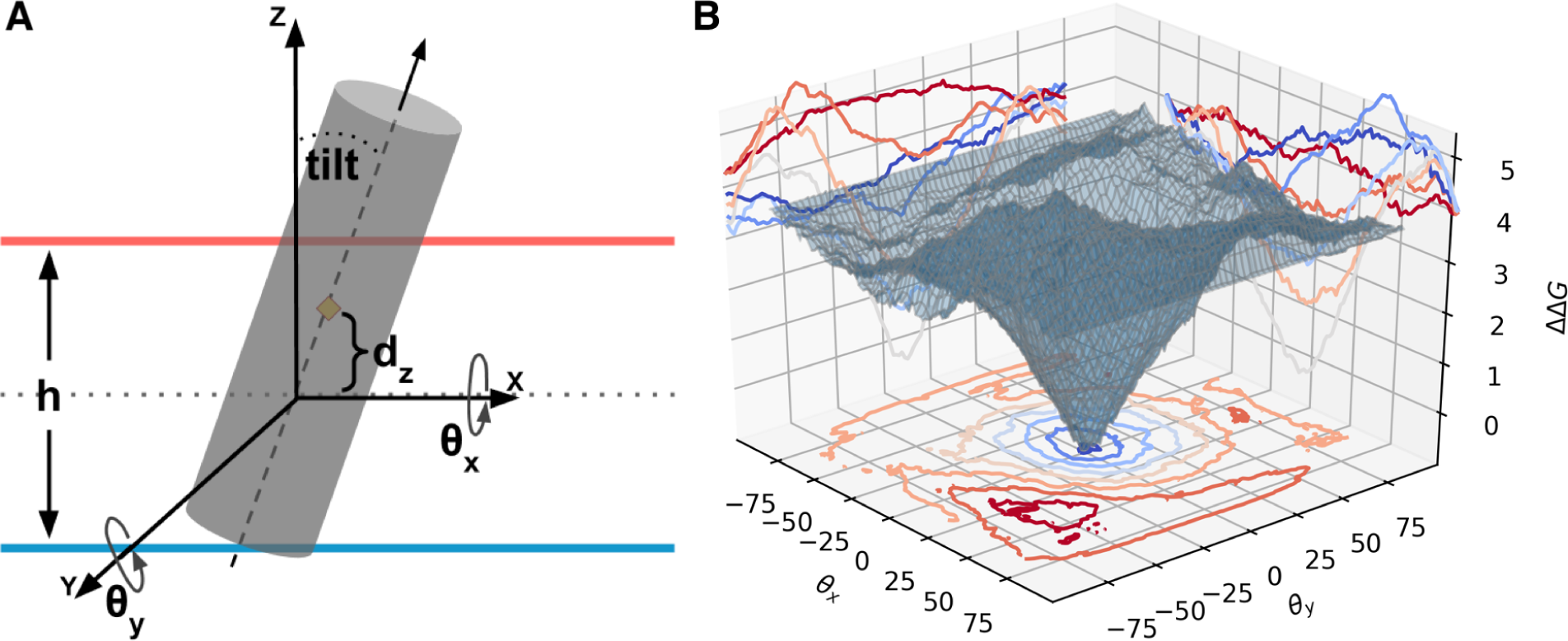
Prediction of positioning and orientation of *β*MPs. **A.** The positioning and orientation of the *β*MP inside the membrane are determine by the rotation angles *θ*_*x*_ and *θ*_*y*_, the translation displacement *d*_*z*_, and the membrane thickness *h*. **B.** The funnel–like landscape of the stability of the BtuB protein. It shows how rotation angles affect the stability of BtuB when *d*_*z*_ and *h* are fixed.

We systematically examine the parameter combination of *θ*_*x*_, *θ*_*y*_, *d*_*z*_, and *h*. A *β*MP is predicted to take the position and the orientation when the lowest free energy is reached. The predicted protein tilt angles of all 58 *β*MPs correlate well (*r* = 0.76) with OPM records.^29^ The average protein tilt angle of 7.3° is consistent with that of 6.2*±*1.8° recorded in the OPM. The strand tilt angles and the membrane thickness predicted are again in good agreement with experimentally determined results (Table 1).

**Table 1.** Comparison between the predicted positioning and orientation and experimental results^31^ of *β*MPs. ^*\**^ The experimentally measured tilt angles are the upper bounds of the actual values.^31^

|  | Protein | PDB ID | Experiment | GeTFEP | OPM |
| --- | --- | --- | --- | --- | --- |
| TM tilt<br>(°) | FhuA | 2fcp | 46.0* | 38.2 | 38.3 |
|  | OmpA | 1bxw | 44.5* | 40.2 | 38.7 |
| Membrane<br>thickness<br>(Å) | FhuA | 1fep | $\geq 23.1$ | 23.5 | 24.3 |
| | OmpF | 2omf | $\sim 21.0$ | 22.8 | 25.2 |
| | BtuB | 1nqe | $\geq 20.2$ | 23.0 | 23.4 |

**Table 2.**
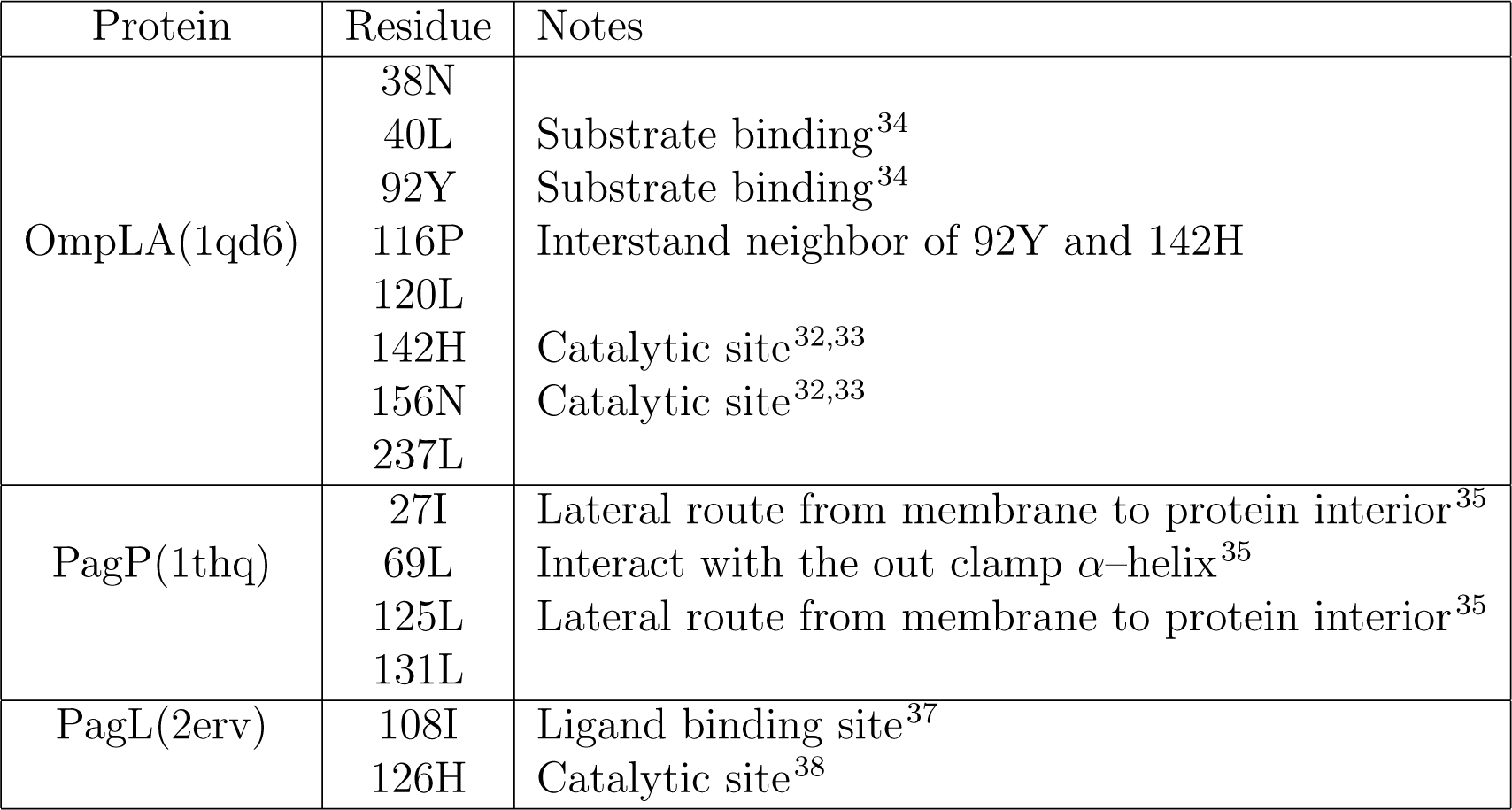
Predicted important sites in OmpLA, PagP and PagL by deviation analysis

### Prediction of structurally and functionally important sites of *β*MPs

While overall the computed TFEs of lipid–facing residues of *β*MPs follow the general pattern of the GeTFEP, the TFE values of a specific residue in a particular *β*MP can deviate significantly from values in the general profile (see SI for details). Among all 3,500 lipid–facing residued in the TM segments of all 58 *β*MPs, we find that 305 or 8.7% of the residues have TFE values deviate significantly from the GeTFEP. Since lipid–facing residues are overall the major contributors to the stability of *β*MPs as discussed above, the deviation from the general profile indicate that the residue is likely to have important roles other than providing stability. To understand the origin of these deviations, we examined three proteins in details, namely, OmpLa, PagP, and PagL, which have sufficient experimental information. We found that most deviant residues either have functional roles or have local structures quite different from residues in the canonical model of beta barrels (Tab 2).

Among the deviant residues in OmpLa, 142H and 156N are both in the catalytic triad^32,33^ that are essential for its phospholipase activities; 40L and 92Y are the sites where substrates bind;^34^ Furthermore, the deviant residue 116P interacts with 92Y and 142H through hydrogen bonds. Among the deviant residues in PagP, 69L interacts with the out–clamp *α*–helix of PagP;^35^ 27I and 125L are both at the lateral routes where *β*–hydrogen bonding is absent (Fig 6), which ensure that substrates can access the protein interior so that PagP can carry out its enzymatic functions.^36^ In PagL, the deviant residue 108I is in the ligand binding site,^37^ and 126H is part of the catalytic triad of its enzymatic site.^38^

**Figure 6.**
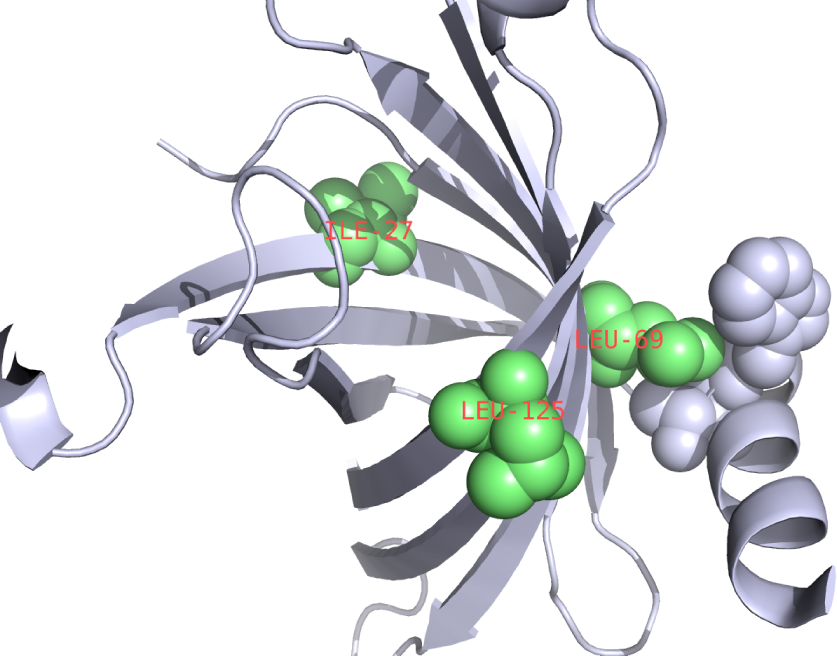
Predicted important sites of PagP. Residues 27I and 125L are at the sites where the hydrogen bonds between the *β*–strands are disrupt. 69L has interaction with the out–clamp *α*–helix of PagP.

As the calculation of TFEs does not require knowledge of 3D structures of *β*MPs, our results suggest that deviation analysis can help to discover functional sites and/or structurally anomalous sites using sequence information only. While our analysis is restricted to three proteins due to the limited nature of experiment data, we believe overall deviant residues play special roles in either performing biological function or in maintaining the unique structural form of *β*MPs.

### GeTFEP can predict TM region of *α*–helical membrane proteins

Although the MF–scale was measured in the *β*MP system, it was suggested that the scale is also applicable to TM region of the *α*–helical membrane proteins (*α*MPs), since the MF–scale has a strong correlation with the nonpolar solvent accessible surface areas of the residues.^10^ We hypothesize that the GeTFEP may also reflects fundamental thermodynamic properties of transferring sidechains of amino acids to the membrane environment, regardless whether the residue is in a *β*–barrel or a *α*–helical membrane protein. We carried out the standard hydropathy analysis^39^ using the Membrane Protein Explorer (MPEx) program.^40^ on 131 *α*MPs obtained from the MPTopo database^41^ Since MPEx uses depth–independent hydrophobicity scales, we used the mid–GeTFEP scale for our calculation.

The results show that this simple analysis using the mid–GeTFEP scale correctly predicts both the TM regions and the numbers of the TM segments for 90 or *∼*69% of the 131 *α*MPs in the dataset. This compares favorably to other hydrophobicity scales, including those measured or derived from *α*MPs (Table 3). For most of the remaining 41 proteins, GeTFEP correctly predicted the TM regions, but predicted the numbers of the TM segments incorrectly due to the ambiguity in assignment of whether two consecutive TM segments should be considered as one TM segment (see Fig. S5B for an example). Examination of the number of TM residues correctly predicted by the mid–GeTFEP scale show that we achieves a precision of *∼*85% and a recall of *∼*71%, which compares favorably to other hydrophobicity scales (Table 3). These results suggest that the GeTFEP reflects fundamental thermodynamic properties of amino acid residues inside membrane, and can be used to study the general stability of both *α*–helical and *β*–barrel membrane proteins.

**Table 3.** Prediction of TM segments and residues *α*MPs. The mid-GeTFEP scale performs better than the other hydrophobicity scales. The first three scales are measured or derived in *α*–helical systems, the others in *β*MPs.

| Hydrophobicity scale | αMPs % (#) with TM segs. correctly predicted | TM res. precision | TM res. recall | TM res. F-measure |
| --- | --- | --- | --- | --- |
| WW-scale | 50%(66) | 73% | 75% | 0.74 |
| Bio-scale | 22%(29) | 95% | 21% | 0.34 |
| $E_Z\alpha$ | 49%(64) | 71% | 77% | 0.74 |
| MF-scale | 48%(63) | 77% | 65% | 0.70 |
| Pro-swapped | 49%(64) | 72% | 74% | 0.71 |
| mid-GeTFEP | 69%(90) | 85% | 71% | 0.78 |

#### The validity of transfer free energy value of Pro in the GeTFEP

We further examine the TFE value of Pro in the mid-GeTFEP scale, which is qualitatively different from that in the MF–scale. We swapped the value of Pro from MF–scale into the mid–GeTFEP scale, and used this Pro–swapped scale in the hydropathy analysis. This is reasonable as the mid–GeTFEP scale is strongly correlated with the MF–scale, and has comparable values. However, we found that the precision of predicting TM residues deteriorates significantly from 85% using the mid–GeTFEP scale to 72% using the Pro–swapped scale (Table 3). This result suggests that Pro is more likely to be membrane unfavorable as characterized by the mid–GeTFEP scale rather than membrane favorable as characterized by the MF–scale.

## 3 Conclusions and discussion

In this study, we derived the General Transfer Free Energy Profile (GeTFEP) from a non– redundant set of 58 *β*MPs. We showed that the GeTFEP agrees well with previous experimentally measured and computationally derived TFEs. The GeTFEP reveals fundamental thermodynamic properties of amino acid residues inside membrane environment, and it is useful in analysis of stability and function of membrane proteins.^5^

As the lipid membrane bilayer is anisotropic along the bilayer normal,^42^ a residue at different depth of the membrane will have different interaction with lipid molecules in the environment, resulting in the depth–dependency of transfer free energies. However, there are few experimental measurements of TFEs at different depth positions other than the hydrophobic core, except Arg and Leu.^10^ Comparison between the GeTFEP and the experimentally measured values of Arg and Leu shows that the GeTFEP captures this depth–dependency well.

In addition, the GeTFEP exhibits asymmetric values between TFEs of residues in the membrane inner leaflet (depth −4 to 0) and in the outer leaflet (depth 0 to +4, Fig. 2C). Most *β*MPs in our dataset resides in the bacterial outer membrane, whose outer leaflet contains additional complex lipolysaccharides in contrast to its inner leaflet of phospholipids. This asymmetry in membrane composition results in the asymmetry of the transfer free energies in the GeTFEP. To understand membrane proteins in an environment of symmetric membrane leaflets, we also derived a symmetric TFE profile, named sym–GeTFEP, by mirroring the TFE values of the inner leaflet side of the GeTFEP (Fig. S6). In this study, the sym– GeTFEP was used to analyze the non–outer–membrane *β*MPs, e.g. *α*– and *γ*–hemolysins and vibrio cholerae cytolysin.

We explored the energetic contribution of different regions of *β*MPs during the membrane insertion process. Our analysis showed that the stability of *β*MPs does not come alone from the extensive property of the hydrophobicity of lipid–facing residues in the TM segment. Rather, the pattern of the amino acid residues in the TM segment also play significant roles. Results from analysis of sequence shuffling show that the patterns and location of amino acid residues are optimized to stabilize *β*MPs in the membrane environment. Using the GeTFEP, we are also able to predict membrane positioning and orientations of *β*MPs.

The GeTFEP can also be used to detect structurally or functionally important residues in *β*MPs. This can be achieved by examination of residues whose TFEs deviate significantly from the GeTFEP. As calculation of TFEs of residues of a specific *β*MP only requires rough estimation of relative positions between adjacent *β*–strands, which can be reliably predicted from the protein sequence,^43,44^ computing the TFE deviation therefore requires only sequence information. The GeTFEP–deviation analysis can aid in discovery of functional sites or structurally important sites in novel *β*MPs, without requiring knowledge of their 3D structures. In addition, GeTFEP–based analysis can aid in design and engineering of novel *β*MPs.

Furthermore, we demonstrated that GeTFEP can be used to predict TM residues of *α*–helical membrane proteins. Results showed that GeTFEP performs better than the hydrophobicity scales measured/calculated in *α*MP systems, suggesting that the GeTFEP reflects fundamental thermodynamic properties of amino acid residues inside membrane, and can be used to study the general stability of both *α*–helical and *β*–barrel membrane proteins.

## Acknowledgement

This work is supported by NIH R01GM079804, R01CA204962-01A1, R21AI126308, and R35GM127084.

## Supplementary information

### Dataset

We use 58 non–homologous *β*–barrel membrane proteins with less than 30% pairwise sequence identity for this study. The PDB IDs are: 1a0s, 1bxw, 1e54, 1ek9, 1fep, 1i78, 1k24, 1kmo, 1nqe, 1p4t, 1prn, 1qd6, 1qj8, 1t16, 1thq, 1tly, 1uyn, 1xkw, 1yc9, 2erv, 2f1c, 2f1t, 2fcp, 2gr8, 2lhf, 2lme, 2mlh, 2mpr, 2o4v, 2omf, 2por, 2qdz, 2vqi, 2wjr, 2ynk, 3aeh, 3bs0, 3csl, 3dwo, 3dzm, 3fid, 3kvn, 3pik, 3rbh, 3rfz, 3syb, 3szv, 3v8x, 3vzt, 4c00, 4e1s, 4gey, 4k3c, 4pr7, 4q35, 7ahl, 3b07, 3o44.

### Clustering of *β*MP TFE profiles

In this study, euclidean distance between the TFE profiles of the *β*MPs and single linkage are used in the hierarchical clustering. The silhouette score is a measure for assessing the quality of clustering results. In practice, a *>* 0.5 silhouette score indicates a good clustering Our results show that it is not reasonable to cluster the *β*MPs into two groups, since the silhouette score is *<* 0.5 at 2 clusters (Fig. S1A). When we increase the number of clusters, the silhouette score keeps decreasing. Therefore, we conclude that only one group exists for our *β*MP dataset. Fig. S1B visualizes the cluster result after the profiles of *β*MPs are reduced to a 3D space using the Principal Component Analysis (PCA).

**Figure S1.**
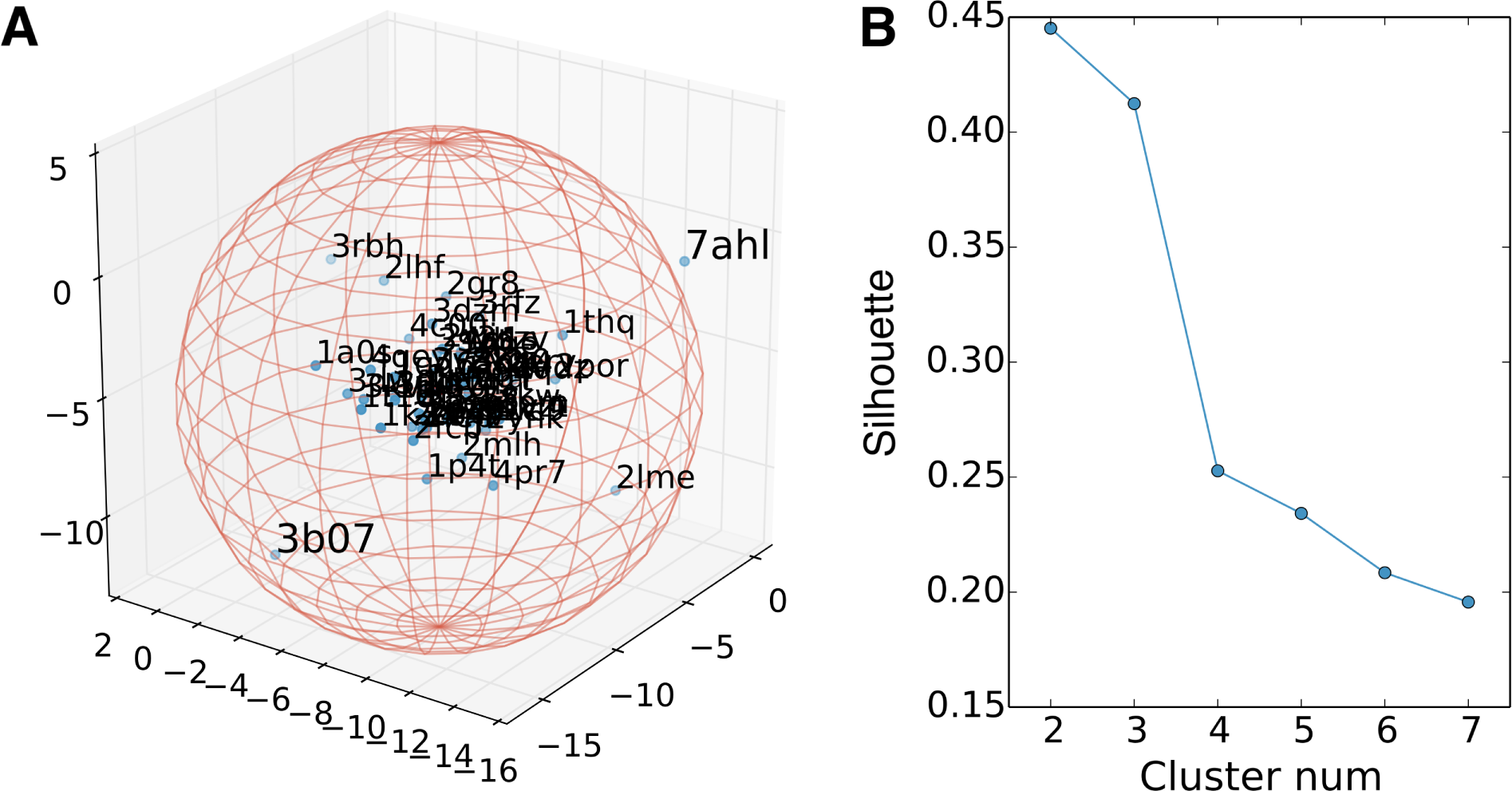
**A.** Visualization of the *β*MP TFE profiles after reduced to a 3D space via PCA. **B.** The silhouette scores for different cluster numbers of the *β*MP TFE profiles.

In the hierarchical clustering, we also tried other parameter settings with correlation distance and/or other reasonable linkages (eg. average linkage or weighted linkage), and the conclusion remains the same.

**Figure S2.**
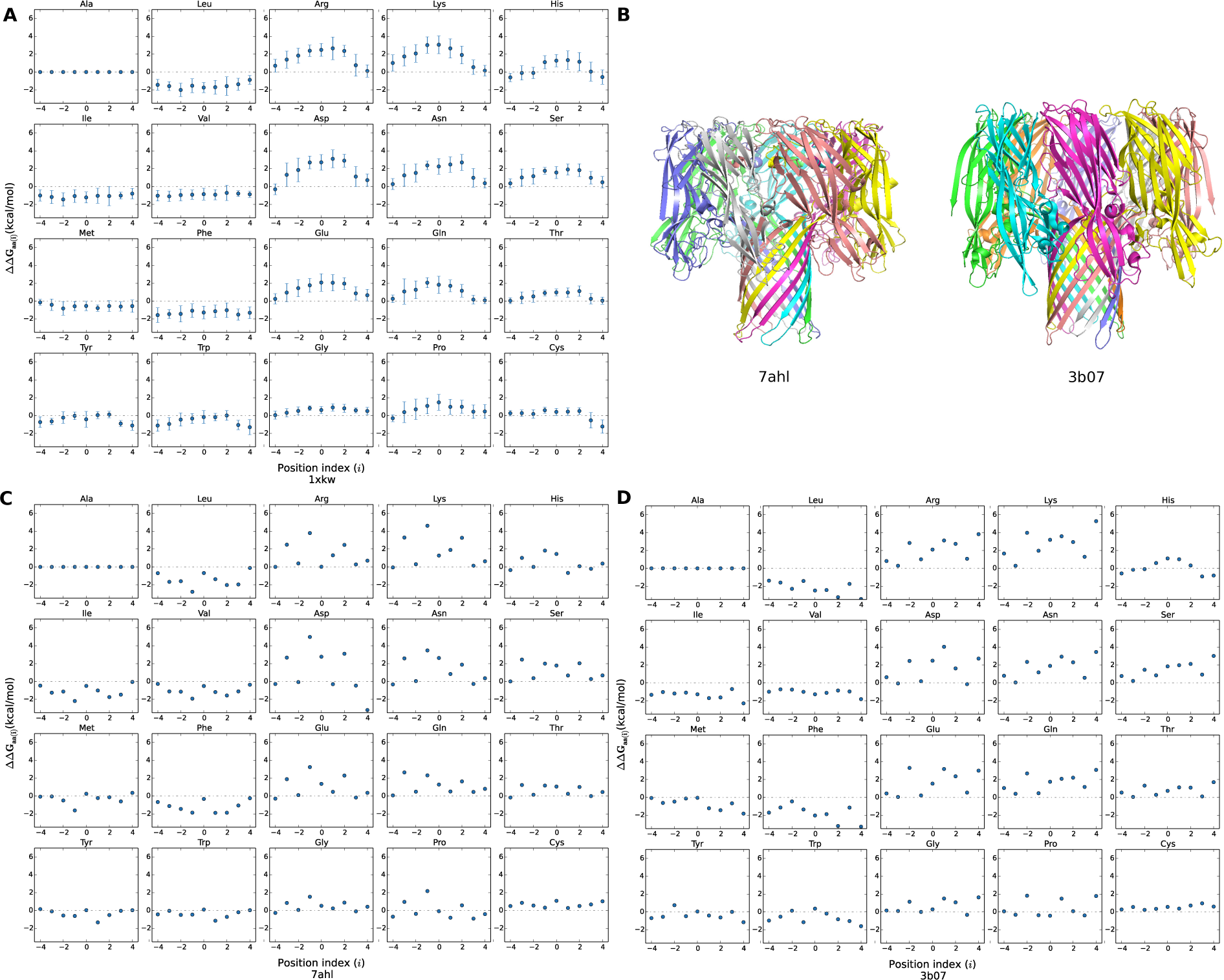
Structures and TFE profiles of *α*– and *γ*–hemolysin. **A.** A typical TFE profile of *β*MPs (FptA, PDB id:1xkw). **B.** The structures of *α*–hemolysin (PDB id:7ahl) and *γ*–hemolysin (PDB id:3b07) Both TM segments are constructed with repeated *β*–hairpins. **C.** The TFE profile of *α*–hemolysin. **D.** The TFE profile of *γ*–hemolysin. Since the structures are both repeated hairpin, there is only one data point for each amino acid residue in every depth of their profiles.

**Figure S3.**
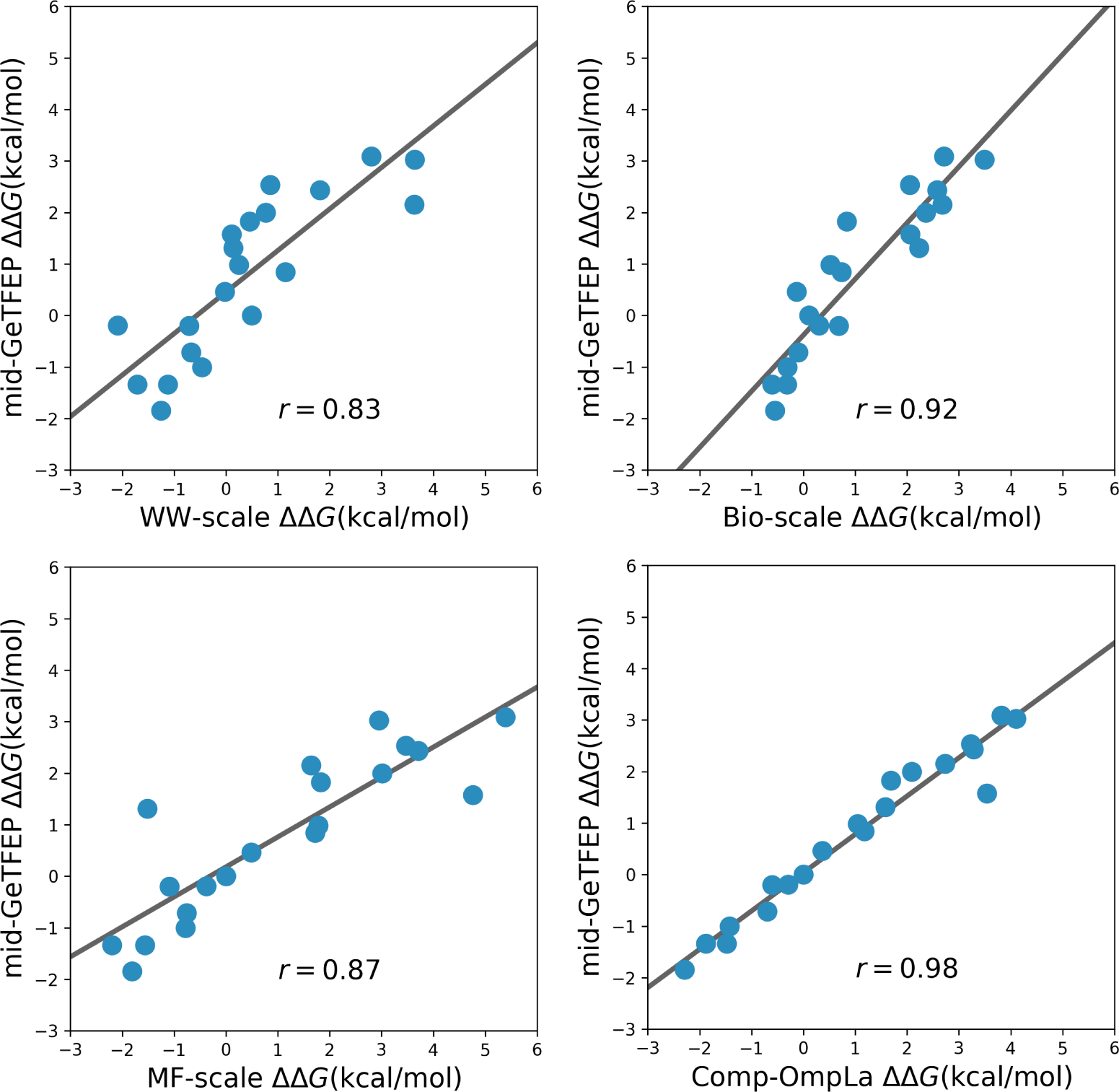
Comparison between the mid–GeTFEP scale and other hydrophobicity scales. The mid-GeTFEP scale agrees well with previously measured or derived hydrophobicity scales.

**Table S1.**
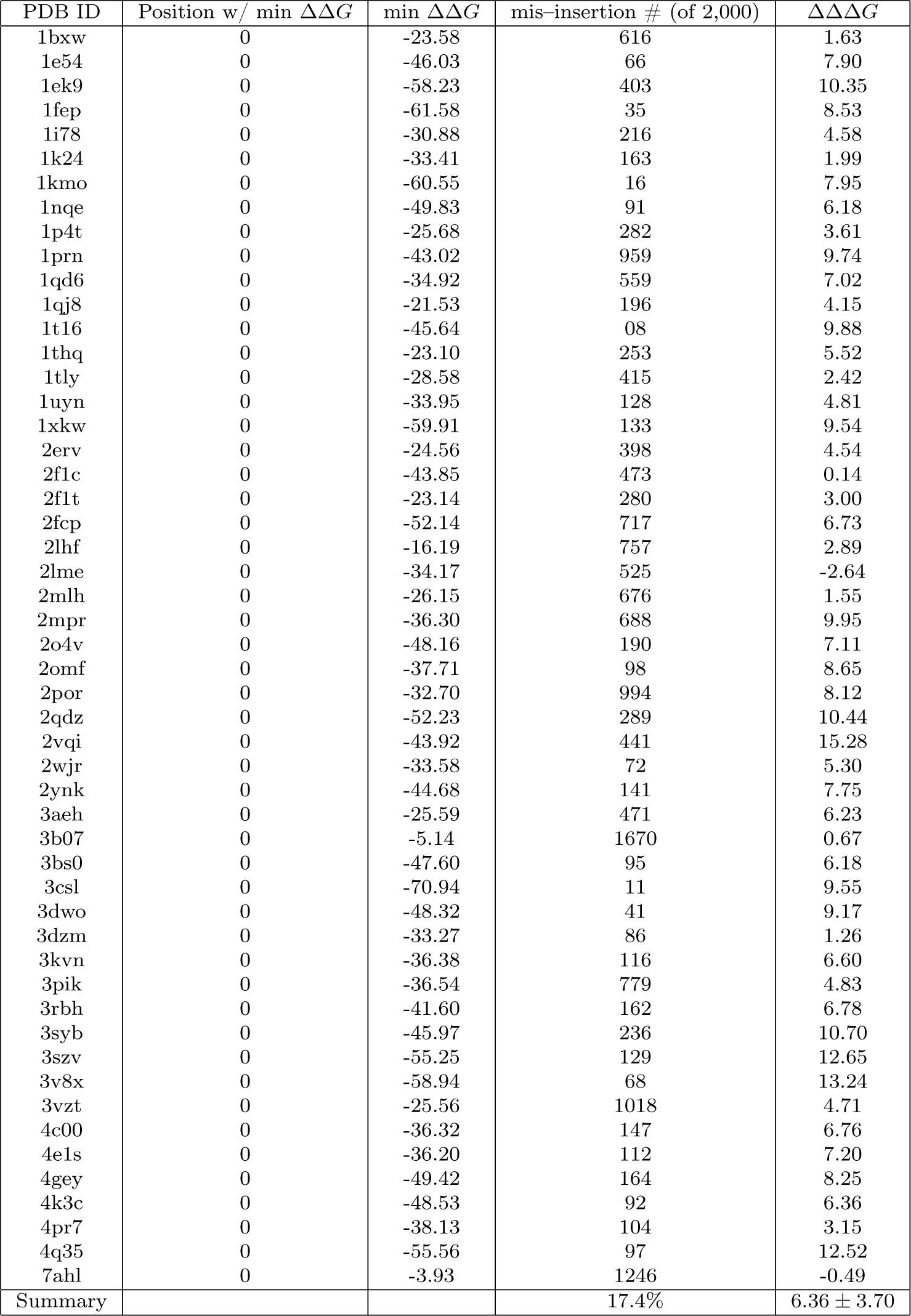
The insertion TFEs of WT and lipid–facing residue–shuffled *β*MPs calculated with GeTFEP. The ΔΔ*G* shows the differences between TFEs of the WT *β*MPs at step 0 and the average of the minimum TFEs of the lipid–facing residue–shuffled *β*MPs.

**Table S2.**
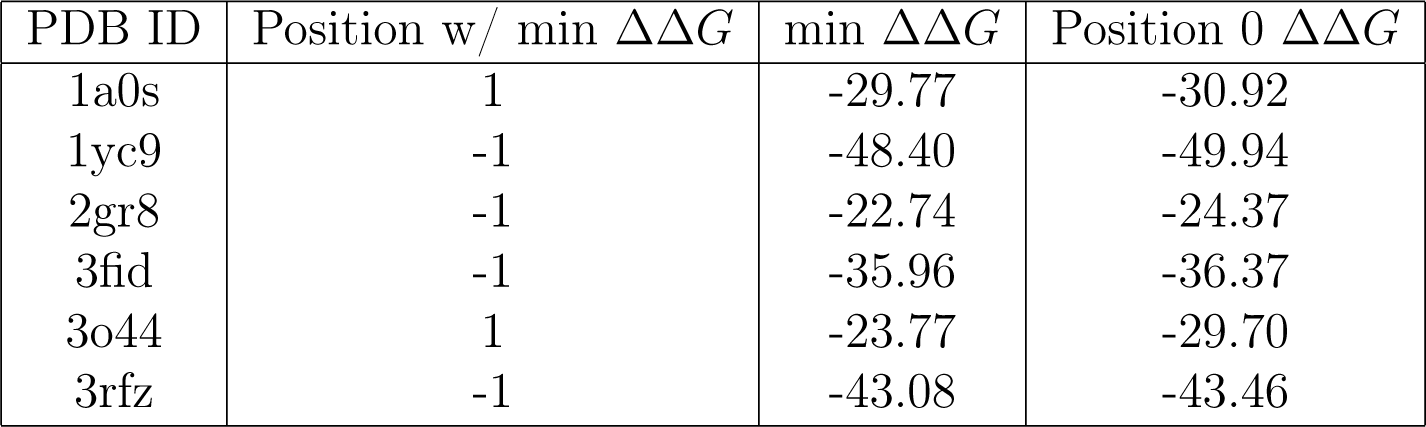
The computed insertion TFEs of the 6 *β*MPs that do not have the minimum energy at position 0. However, their most stable position is close to 0, and the minimum TFEs are close to their TFEs at position 0. Nonetheless, we exclude these 6 *β*MPs in our other analysis of membrane insertion stability.

**Figure S4.**
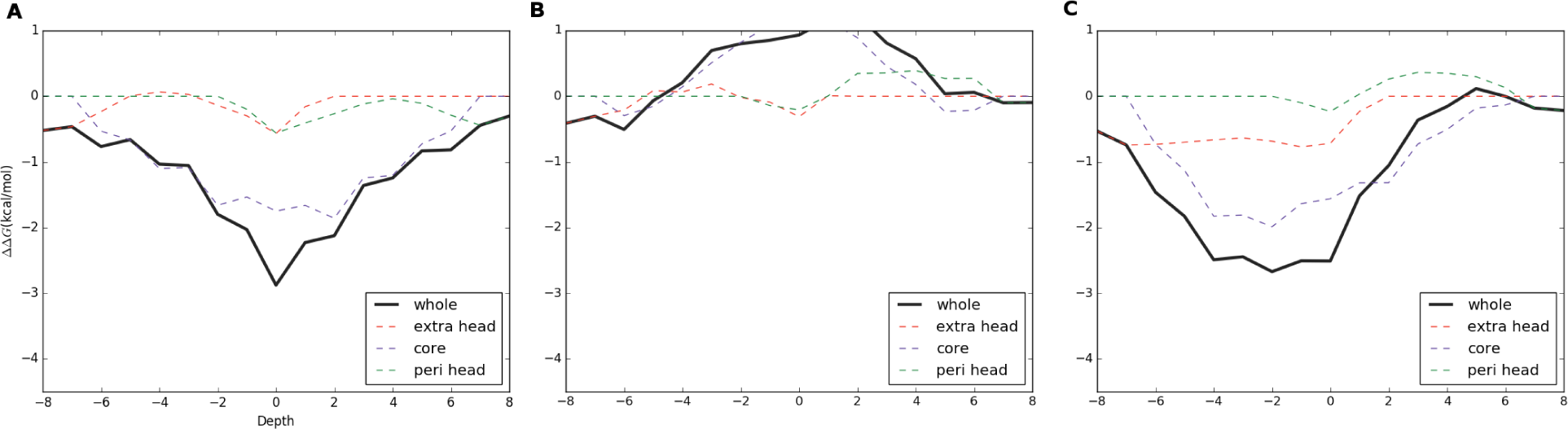
An example of the insertion TFEs (Omp32, PDB ID:1e54). **A.** The insertion TFEs of the intact Omp32 shows a funnel pattern. **B.** The insertion TFEs of the residue–shuffled Omp32 regardless of side–chain directions. **C.** The insertion TFEs of the lipid–facing–residue–shuffled Omp32

### Prediction of structurally or functionally important sites

For a lipid–facing residue in a *β*MP, we calculate the z–score of its TFE by 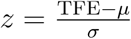, where *µ* and *σ* are respectively the mean and the standard deviation values in GeTFEP of the same amino acid in the same depth. When *z >* 1.64 or *<* −1.64 (which correspond to 5% and 95% in the normal distribution), we take the residue as a deviant that may be structurally and functionally important.

**Figure S5.**
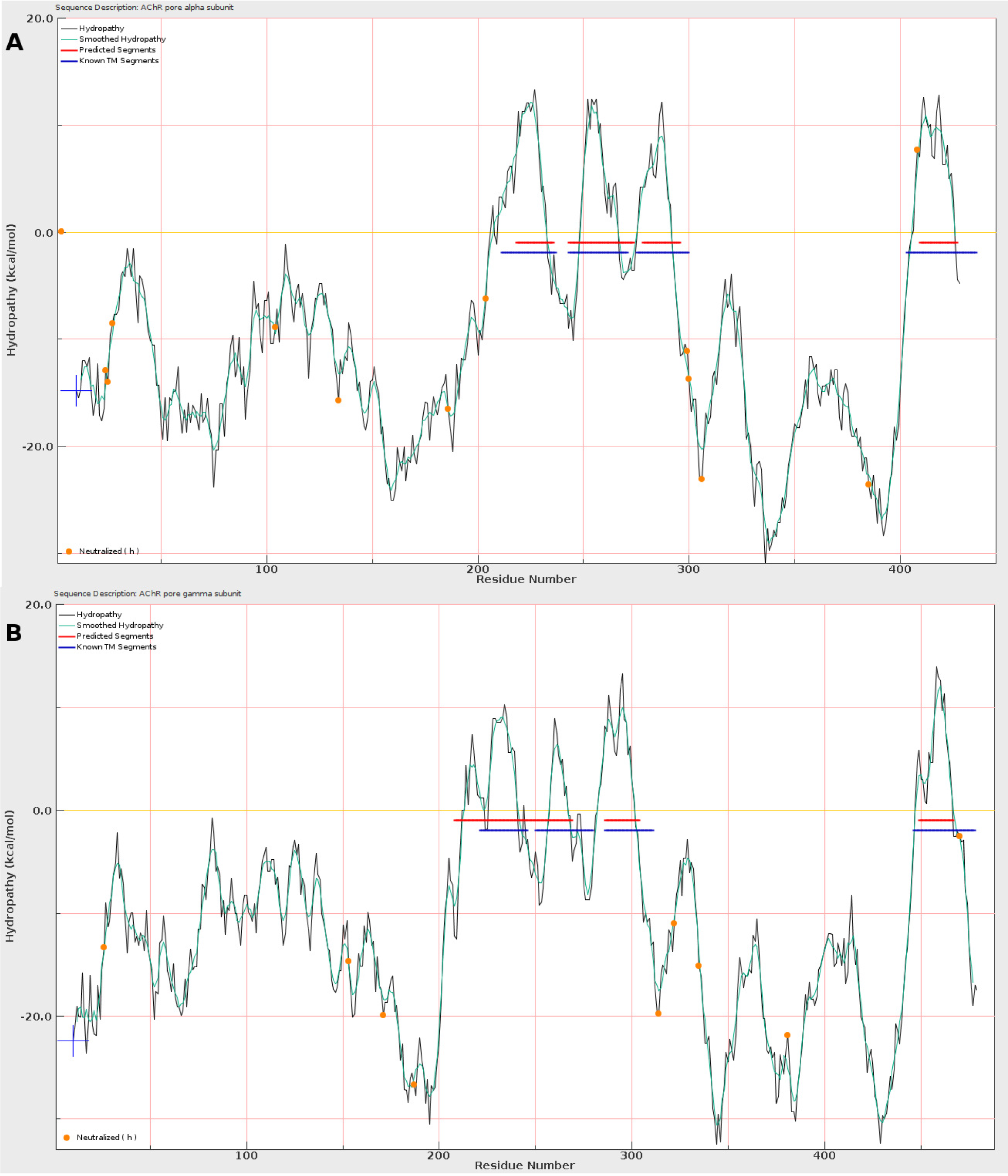
Hydropathy analysis with mid–GeTFEP. The blue segments are the known TM segments, while the red ones are predicted by the hydropathy analysis. The analysis was carried out using Membrane Protein Explorer (MPEx)^40^ **A.** An example (AChR pore *α* subunit) shows both the TM region and the number of the TM segments are correctly predicted. **B.** An example (AChR pore *γ* subunit) shows the predicted number of the TM segments are wrong, though the TM regions are correctly predicted.

**Figure S6.**
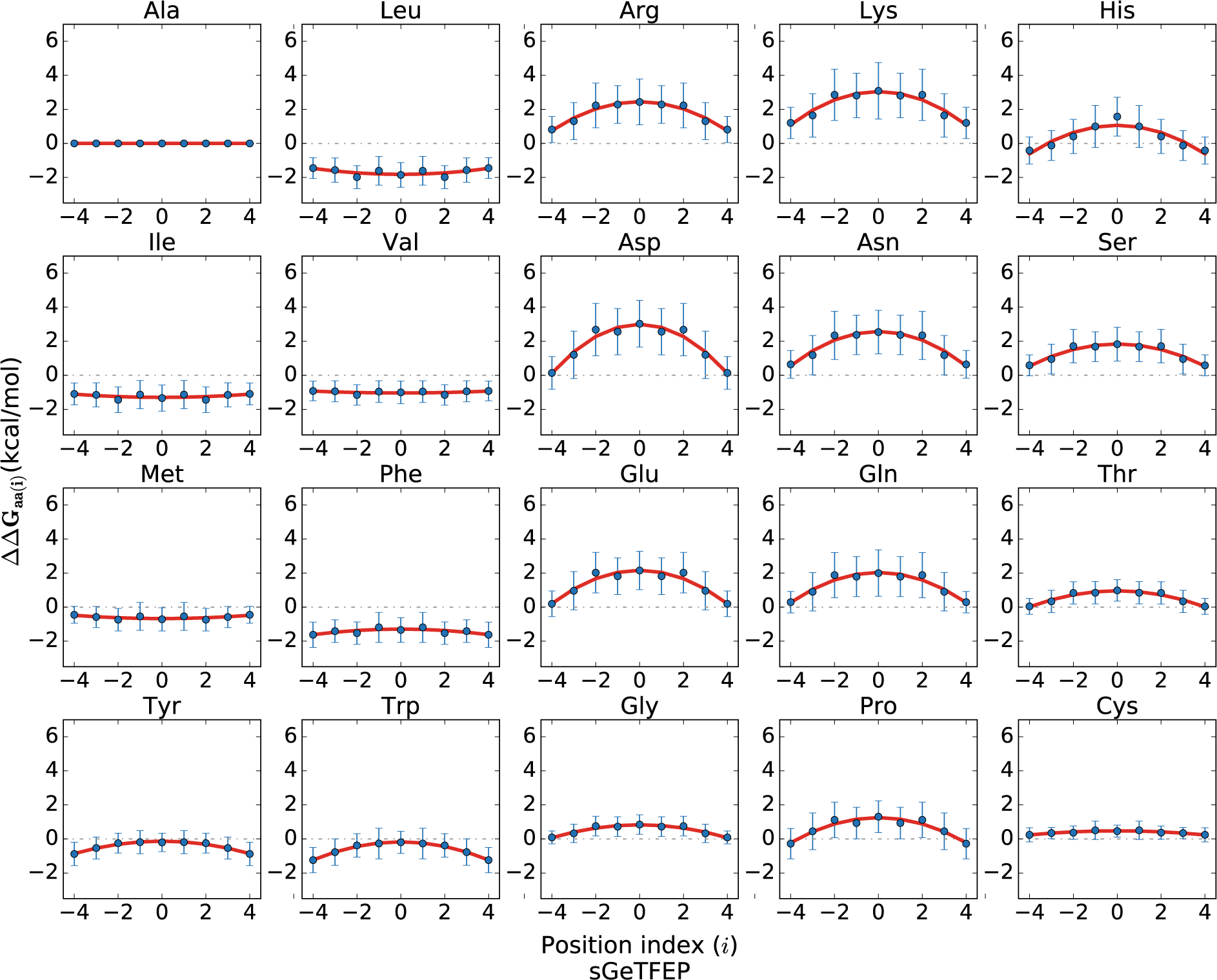
The sym–GeTFEP for symmetric membranes. This profile is derived by mirroring the left part (depth −4 to −1) of the original GeTFEP.

